# Genome and methylome of the root crop table beet (*Beta vulgaris*)

**DOI:** 10.64898/2026.09.07.749830

**Authors:** Thomas Holzweber, Kevin M. Dorn, Laide A. Rasaki, Juliane C. Dohm, Heinz Himmelbauer

**Author notes:** NC State University, Raleigh, NC 27607, USA.

## Abstract

Table beet (*Beta vulgaris subsp. vulgaris var. vulgaris*), also known as red beet, garden beet, or beetroot, is an important crop with both nutritional and agricultural value, yet high-quality genomic resources for this taxon are limited. We present the chromosome-scale genome assembly RefTab-1 based on PacBio and Hi-C data of the inbred table beet genotype W357B. The assembly has a size of 668.6 Mbp and an N50 of 72 Mbp, with 96% of the assembly anchored to nine pseudo-chromosomes. We annotated 64% repetitive DNA, 24,863 protein-coding genes, and 10,405 non-coding RNA loci including a 5S rRNA cluster on chromosome 4. Genome-wide methylation profiling using EM-seq revealed high levels of cytosine methylation in repeats and context-specific methylation in gene bodies and promoters. We propose three terpene synthase candidate genes based on functional annotation and based on their location near a genetic marker on chromosome 8 previously associated with geosmin concentration, responsible for the earthy aroma of table beet. Comparative analysis with sugar beet indicated overall structural conservation and a 9 Mbp inversion on chromosome 9. These high-quality resources for table beet will enhance genetic, epigenetic, and comparative studies in *Beta vulgaris,* advancing beet biology and breeding programs.

## Background

Table beet (*Beta vulgaris* subsp. *vulgaris* var. *vulgaris*), also called red beet, beetroot, or garden beet, is a biennial vegetable crop valued for its pigmented, fleshy taproots (Fig. 1) [1]. Taxonomically, the species *Beta vulgaris* comprises all cultivated beets within the subspecies *vulgaris*, which in addition to table beet includes other economically important crops, i.e. sugar beet, fodder beet, and leaf beets (chard). All these cultivated forms trace their origin back to a common ancestor, the wild sea beet (*Beta vulgaris* subsp. *maritima*), a coastal plant native to the Mediterranean basin and the European Atlantic region [2]. Genetic evidence points towards the Eastern Mediterranean region as the place where beets were first domesticated [3,4].

**Figure 1.**
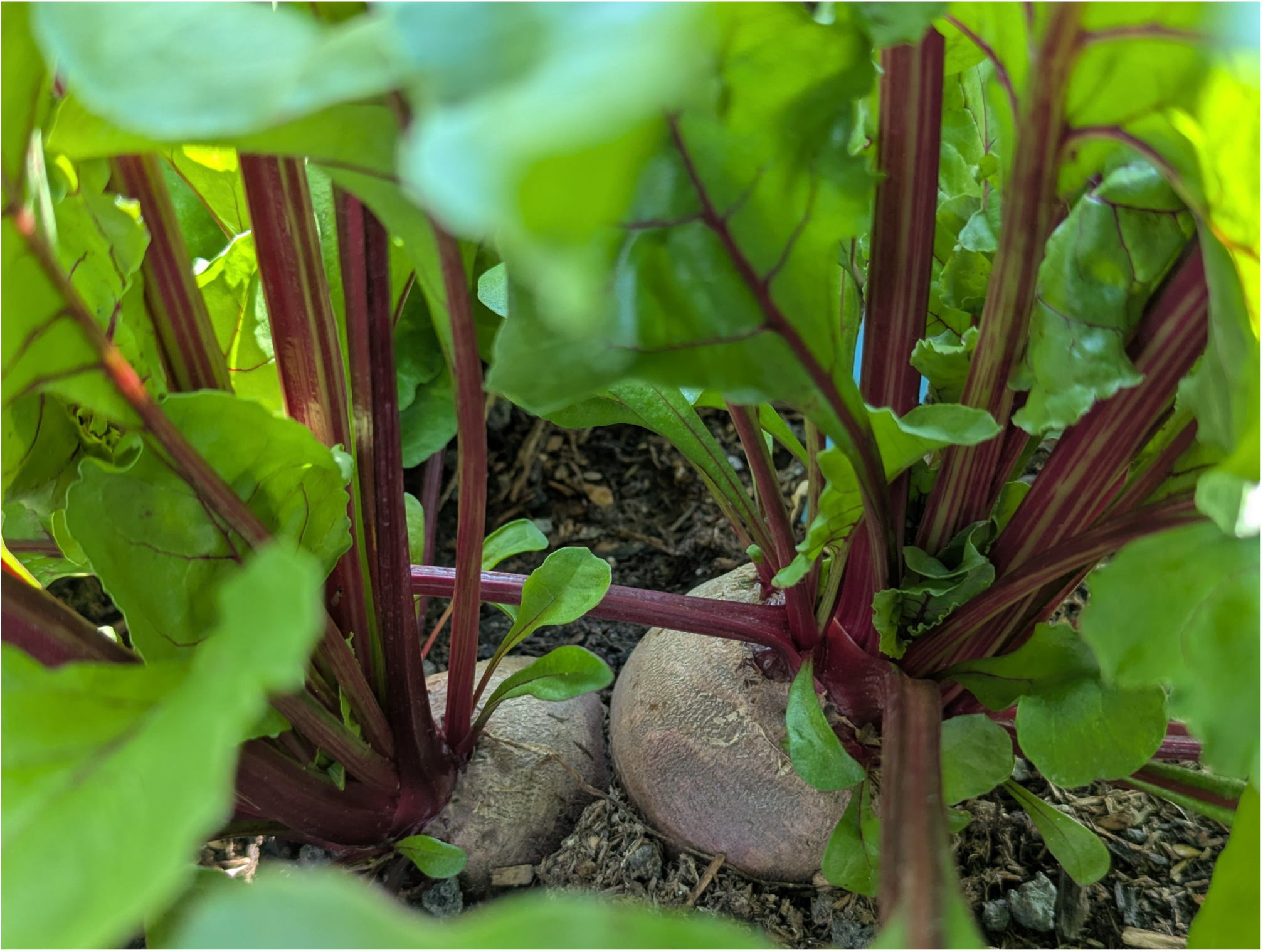
Table beet “Rote Kugel 2” at BOKU botanical garden. Seeds were obtained from Gartenland GmbH (Aschersleben, Germany). The setup was arranged in the context of the Horizon-Europe project “Pro-Wild: Protect and promote crop wild relatives”. Picture taken on June 12th, 2025 and kindly provided by Lukas Schönmann.

Early wild forms of *Beta vulgaris* had slender, often forked roots that were used medicinally, while the leaves were the primary parts that were consumed. The swollen root of the modern beet developed from a feature already present in its wild ancestor, extra growth layers called supernumerary cambia [5]. Through centuries of human selection, this feature was selected, leading to the enlarged, fleshy taproots of table beet, fodder beet, and sugar beet. Historical records suggest that swollen-root beets existed by the 16th century. Today, the appearance of table beet varieties is still diverse, root shapes vary from conical and cylindrical to the familiar spherical form and colors range from red to yellow [1].

Culinary uses of the crop are equally diverse. Raw table beets are healthy and colourful additions to juices, while cooked beets can be added to salads and are an essential constituent of Northern European labskaus and Eastern European borscht. The table beets’ distinctive flavor is mainly defined by sweetness and earthiness. The sweet taste is attributed to a high sucrose content, while the earthy aroma comes from a terpenoid called geosmin [1]. A locus on chromosome (BvChr) 8 has been associated with geosmin concentration in table beets [6]. While considered a minor crop globally, it still has substantial regional economic importance. Poland, France, and Germany are the top three producers in the European Union [7]. In the United States, the largest shares of table beet acreage are found in New York, Wisconsin, and California [8].

Because of its economic importance, sugar beet has historically been the primary focus of genomic research within the genus. This has resulted in a series of reference genomes [9–14] and other molecular resources for sugar beet [15–19]. Further relevant resources to study beets include reference genomes for chard [20] and the wild beet taxa sea beet and *Beta patula* [21,22]. In table beet, earlier work has developed a first-generation genetic map from a sugar beet × table beet cross [23]. Several studies have generated re-sequencing data from table beet accessions, either from pooled individuals or from single plants [24,25]. At the trait level, transcriptome sequencing mapped the table beet *R* locus (responsible for red versus yellow pigmentation) to the *CYP76AD1* gene, serving as a reference point for tracing the genetic divergence of pigment pathways in related Caryophyllales crops [26,27]. A preliminary sequence assembly for the table beet inbred W357B was generated to support read alignment and QTL mapping [28,29]. While this resource provided a structural backbone, it was based on a reduced set of sequencing data and lacked gene prediction or additional genomic features like repeat annotation and epigenomic information.

Aside from the primary DNA sequence, chemical modifications to nucleotides are crucial for regulation of genome activity. One of the most studied types of epigenetic modification in plants is cytosine methylation (5mC). In plants, this methylation occurs mainly in CG, CHG, and CHH contexts (where H can be A, T, or C) [30]. DNA methylation is essential for genome stability through silencing of transposable elements and other repetitive sequences [31]. It also plays a critical role in controlling gene expression. Methylation of promoter regions can alter their accessibility and therefore prevent binding of transcription factors and RNA polymerase [32]. Methylation of specific promoter regions can be dynamic in response to environmental factors like drought, temperature, or nutrient stress, enabling adaptation to various circumstances and fine-tuned regulation of stress-response genes [33]. CG methylation, mainly maintained by DNA methyltransferase 1 (MET1), is often found in gene bodies and is associated with constitutive gene expression [34]. CHG methylation, in contrast, is maintained by chromomethylase 3 (CMT3) and plays an important role in silencing transposons in heterochromatin [35]. CG and CHG methylation are both symmetric, meaning both strands of DNA have complementary cytosines that can be methylated and through this, methylation can be conserved during replication. In contrast, CHH methylation lacks a complementary methylated cytosine on the opposite strand and is therefore asymmetric, requiring *de novo* establishment after each replication cycle [36]. CHH methylation is crucial for silencing of transposable elements in euchromatic regions and can be very dynamic in response to environmental stress [37].

DNA methylation has previously been investigated in sugar beet and in broader comparative plant methylome studies. In sugar beet, genome-wide bisulfite sequencing of leaves and leaf-derived callus showed that cytosine methylation followed the typical plant pattern CG > CHG > CHH, was enriched in repetitive sequences, and differed between leaves and callus, particularly in transposable elements and genes [38]. In a comparative analysis of 34 angiosperm methylomes, *Beta vulgaris* was also identified as a highly methylated species across CG, CHG, and CHH contexts [39]. These studies indicate that *Beta vulgaris* genomes are characterized by high levels of cytosine methylation, especially in repeat-rich regions. However, these studies focused on sugar beet or used broad comparative datasets, and a table beet-specific methylome has not yet been described.

Considering the agricultural importance of table beet as a versatile crop used in food production, we set out to provide an annotated, chromosome-scale assembly for this beet crop type. Specifically, we take advantage of the inbred table beet line W357B [40] to generate a table beet reference genome sequence called RefTab-1. Based on the RefTab-1 genome assembly, we assess the table beet repeat landscape, predict both protein-coding genes and non-coding RNAs, analyze genome-wide cytosine methylation patterns, and compare the genomes of table beet and sugar beet to explore structural similarities and differences between these taxa.

## Results

### Table beet genome assembly and quality assessment

The genome of the inbred table beet line W357B was sequenced on the Pacific Biosciences (PacBio) Sequel II platform, independently by the USDA and by BOKU University. The resulting datasets were combined, yielding 2.76 million High-Fidelity (HiFi) reads with an average read length of 11,800 bp after correction. The initial assembly of these HiFi reads comprised 2,336 contigs with a total size of 748.7 Mbp and an N50 size of 5.2 Mbp, representing an approximate two-fold increase in contiguity compared to assemblies based on individual datasets. A contamination scan identified 73 contigs, totaling 1.7 Mbp, that fully matched the genome of the plant-feeding mite *Tetranychus urticae*. These contigs were removed (Table 1); the resulting assembly was called RefTab-0.

**Table 1.**
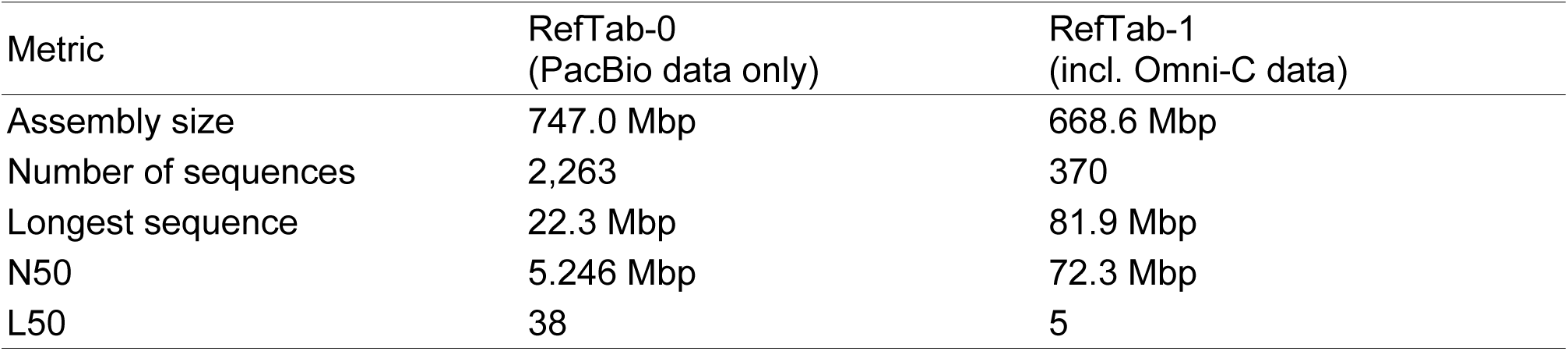
Assembly statistics of RefTab genome assemblies. The assembly size refers to the total length of all sequences. N50: the length at which 50% of the assembly is contained in sequences of this size or longer; L50: the minimum number of sequences required to cover 50% of the assembly.

In a next step, Hi-C data were used for scaffolding the PacBio-based table beet genome assembly. A total of 151 million quality-filtered Omni-C read pairs were mapped to RefTab-0, and subsequent scaffolding resulted in a more than ten-fold increase in N50 size. The nine largest scaffolds were oriented and chromosomally assigned according to the sugar beet genome assembly RefBeet-3.0 [10]. As additional validation, a comparison with another chromosome-scale sugar beet assembly, FC309 [14], was performed, confirming the chromosomal assignments and orientations derived from RefBeet-3.0. Similar to sugar beet chromosome size trends, the table beet scaffold assigned to BvChr 6 was the largest (81.9 Mbp), while the smallest table beet scaffold corresponded to BvChr 9 (62.1 Mbp).

Chloroplast and mitochondrial genome sequences were identified in a total of 1,626 contigs with lengths ranging from 8 kbp to 826 kbp. These contigs were removed, reducing the assembly size to 675.2 Mbp. In addition, we used the HiFi reads to assemble the table beet chloroplast genome *de novo*. The *de novo* assembled table beet chloroplast was 100% identical to the published sugar beet chloroplast genome (GenBank: NC_059012.1), confirming earlier findings in which different *Beta vulgaris* cultivars (including table beet, sugar beet, fodder beet, and others) shared identical chloroplast DNA restriction fragment patterns [41].

A total of 426 scaffolds (31.9 Mbp) remained unassigned. These were aligned back to the nine chromosome-sized table beet scaffolds to check for duplicated sequences. For 65 sequences, >99% of their length aligned to chromosome-sized scaffolds; thus, they were considered duplicates and were removed. Of the remaining unassigned scaffolds, the largest was 808 kbp and the smallest was 10 kbp. The resulting chromosome-scale table beet assembly had an assembly size of 668.6 Mbp in 370 sequences with an N50 size of 72.3 Mbp (Table 1) and is hereafter referred to as RefTab-1.

To confirm the completeness of RefTab-1, we performed an analysis using the Benchmarking Universal Single-Copy Orthologs (BUSCO) tool with gene sets from both the general eukaryota_odb12 lineage and the more specific eudicotyledons_odb12 lineage. For RefTab-1, we identified 100% complete BUSCOs using the eukaryota dataset and 97.4% complete BUSCOs using the eudicotyledons dataset (Table 2).

**Table 2.**
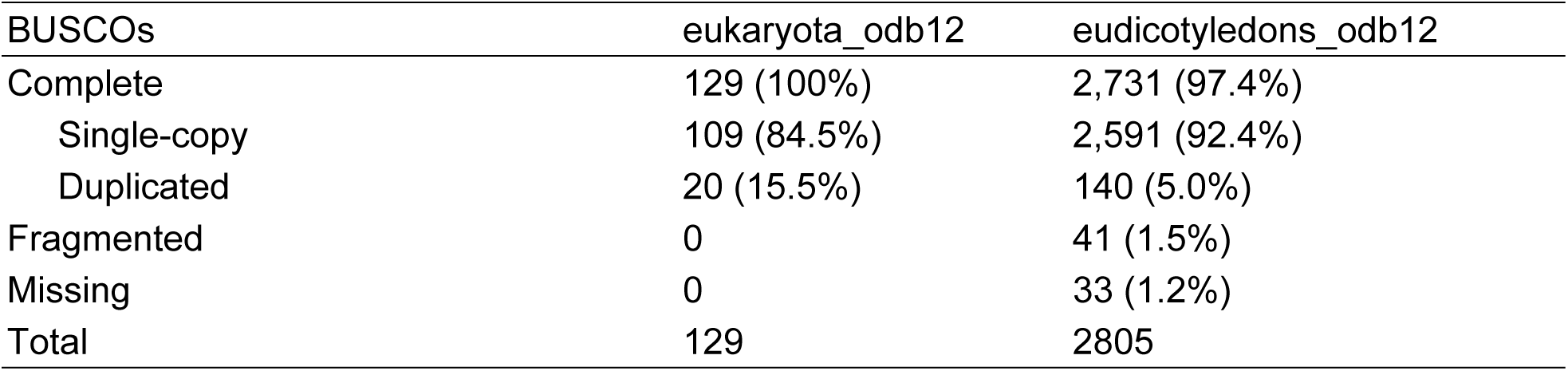
Assessment of genome assembly completeness of RefTab-1. BUSCO analyses were performed with gene sets from both the general eukaryota_odb12 lineage and the more specific eudicotyledons_odb12 lineage.

| BUSCOs | eukaryota_odb12 | eudicotyledons_odb12 |
| --- | --- | --- |
| Complete | 129 (100%) | 2,731 (97.4%) |
| Single-copy | 109 (84.5%) | 2,591 (92.4%) |
| Duplicated | 20 (15.5%) | 140 (5.0%) |
| Fragmented | 0 | 41 (1.5%) |
| Missing | 0 | 33 (1.2%) |
| Total | 129 | 2805 |

Telomeric repeat regions were identified in the chromosomally assigned scaffolds. These repeats, typically consisting of short tandem motifs that protect chromosome ends from degradation and prevent fusion with other chromosomes, were found at the start of all but one chromosome (BvChr 3) and at the end of all but one other chromosome (BvChr 1), indicating accurate identification of chromosomal boundaries in nearly all scaffolds assigned to chromosomes (Fig. 2, Suppl. Fig. S1).

**Figure 2.**
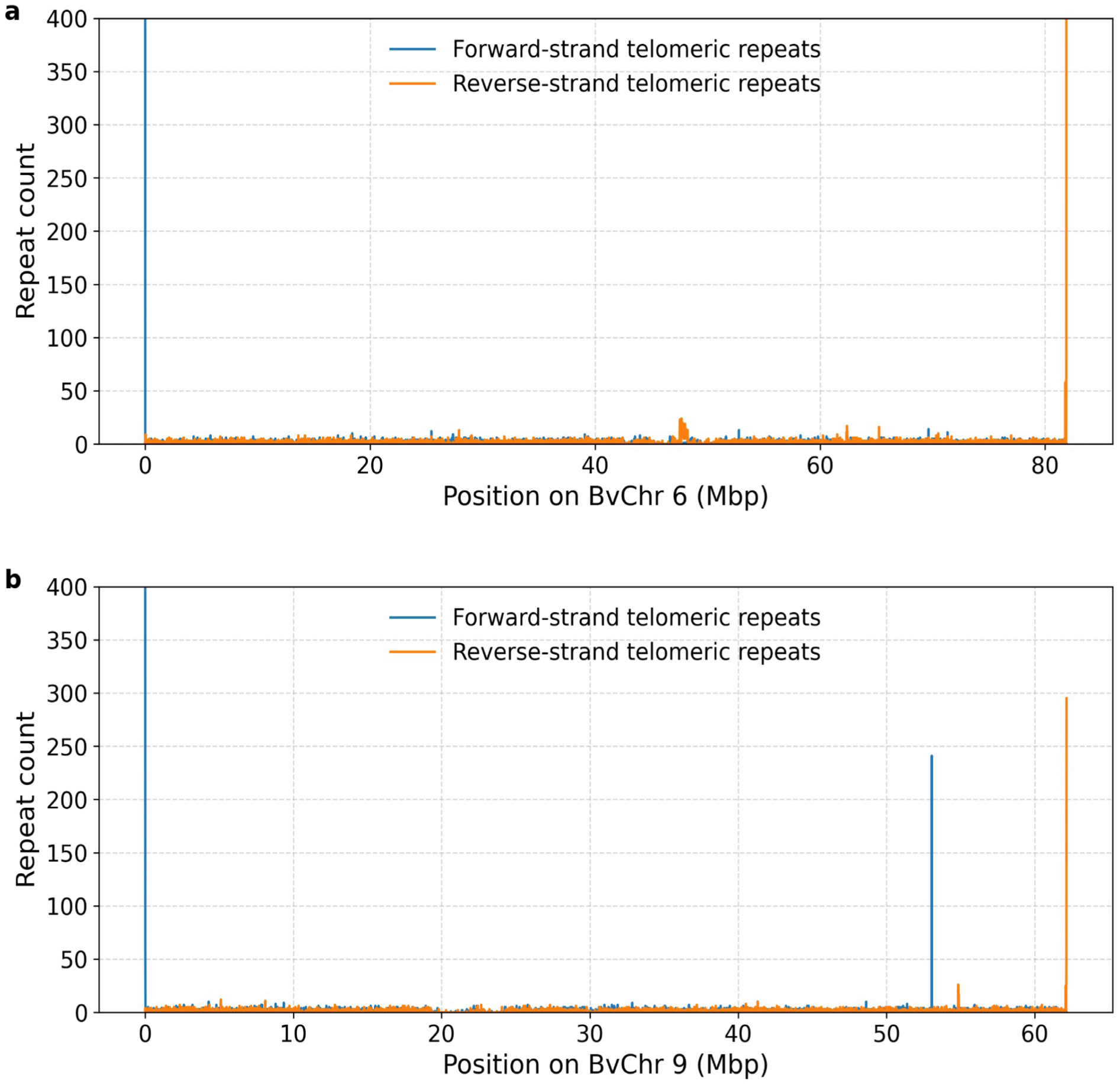
Telomeric repeat sequences detected in BvChr 6 and BvChr 9. Repeat counts represent the number of telomeric repeats identified within 10 kbp sliding windows along each chromosomally assigned scaffold. Blue lines: forward-strand telomeric repeats; orange lines: reverse-strand telomeric repeats. (a) BvChr 6. Both the forward-strand telomeric repeats at the chromosome start and the reverse-strand telomeric repeats at the chromosome end were detected; (b) BvChr 9. Both the forward-strand telomeric repeats at the chromosome start and the reverse-strand telomeric repeats at the chromosome end were detected. Additional forward-strand telomeric repeats were found at a distance of approximately 9 Mbp from the chromosome end.

### Repeat landscape of the table beet genome

To identify and classify repetitive elements within RefTab-1, we employed a two-step approach, starting with repeat modeling followed by repeat masking. RepeatModeler was used with both the Dfam plant database and the Repbase RepeatMasker library, resulting in the identification of 3,280 repeat models, of which 1,683 could be classified. The majority (88% or 1,479) of the classified models were found to be long terminal repeats (LTRs). Long interspersed nuclear elements (LINEs) and DNA transposons were present in equal proportions, forming the second-largest fractions (Table 3).

**Table 3.**
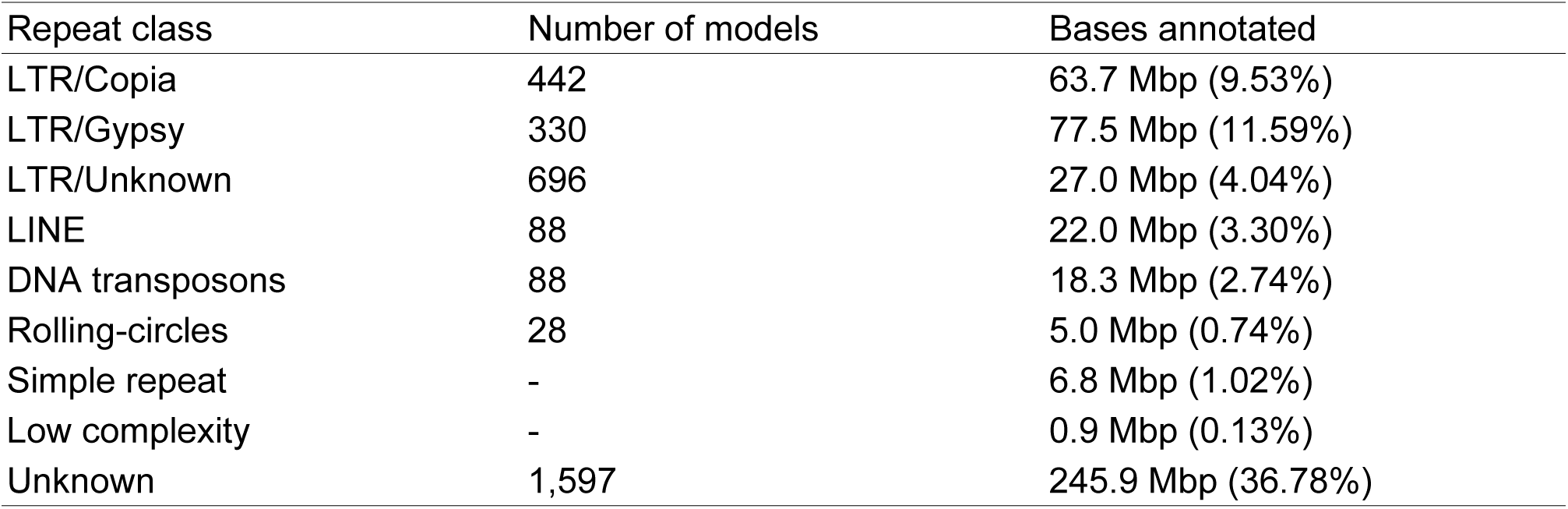
Genome-wide classification and annotation of repetitive elements in RefTab-1. The number of repeat models assigned per class and the total number of bases masked in RefTab-1 for each repeat class. In total, 69.87% of the genome was identified as repetitive. LTR: long terminal repeat; LINE: long interspersed nuclear element.

These repeat models were then used to annotate the genome assembly using RepeatMasker, which categorized 69.87% of the genome as repetitive and was used for repeat masking during gene annotation. The largest portion of these repeats (36.78% of the total assembly size) remained unclassified, while 25.16% were classified as LTR elements, 3.30% as LINEs, and 2.74% as DNA transposons (Table 3).

Additionally, the Extensive *de novo* TE Annotator (EDTA) was utilized to create another, more comprehensive repeat annotation for the final repeat-landscape statistics. The EDTA analysis was supplemented with the RepeatModeler output and the BeetRepeats library as additional inputs. As EDTA was designed to filter out possible false discoveries, the total fraction of the genome classified as repetitive was reduced to 63.68%; however, only 4.57% remained as unclassified, compared to 36.78% when using only RepeatMasker. Approximately 30.63% of the assembly was classified as LTR elements, 11.99% as terminal inverted repeats (TIRs), 8.61% as satellite DNA, and 3.80% as LINEs (Table S1).

### Gene annotation

Protein-coding genes in RefTab-1 were predicted using an integrative approach combining *ab initio* gene prediction, transcript evidence, and homology-based information. To avoid predicting false positives from repetitive DNA, while still preserving useful sequence information, the genome assembly was soft-masked for repeats. Approximately 2.15 billion mRNA-seq read pairs and 408 million single-end reads derived from different sugar beet tissues and developmental stages (leaf, root, seed, seedlings, inflorescence), as well as from stressed plants, were mapped to the assembly. For protein evidence, 5.3 million sequences from OrthoDB Plants v11 were included. Gene models were generated using BRAKER3, which uses RNA-seq alignments, assembled transcripts, and protein homology to produce predictions, and only transcripts with 100% intron evidence were retained.

This approach resulted in a total of 24,863 protein-coding genes (28,672 transcripts), with an average coding sequence length of 1,199 bp and an average of 5.1 exons per gene (Table S2). The largest gene, tab_g17232 on BvChr 7, encompassing nearly 77 kbp, displayed >99% sequence identity to “hypothetical protein BVRB_7g162530” from sugar beet (NCBI GenBank: KMT06182.1) [9]. The table beet gene with the largest coding sequence (CDS), tab_g18400 on BvChr 7, encoding a protein of 5,438 amino acids, was identified as midasin (MDN1), a large and highly conserved protein in eukaryotes, critical for assembly and maturation of the large (60S) ribosomal subunit. There were 35 genes with a CDS length of less than 25 amino acids, 793 genes with a CDS length between 25 and 50 amino acids and 2,075 genes with a CDS length between 50 and 100 amino acids. Genes with a CDS length <100 amino acids are frequently enriched in long non-coding RNAs (lncRNAs); however, some such genes may also encode functional peptides. The resulting gene annotation is hereafter referred to as TabSet-1.

Functional annotation was performed using eggNOG, enabling the assignment of orthologous groups (COG), Gene Ontology (GO) terms, KEGG pathway information (KO), and Pfam domains. Overall, 20,925 (84.2%) of the predicted genes were assigned to a functional category, providing a comprehensive view of the table beet gene repertoire (Table 4, Fig. 3).

**Table 4.** Number of TabSet-1 genes with functional information. The numbers indicate how many genes or transcripts received at least one annotation from the respective databases: COG (Clusters of Orthologous Groups), GO (Gene Ontology), KO (KEGG Orthology), and Pfam (Protein families database).

|  | Genes | Transcripts |
| --- | --- | --- |
| COG | 14,063 | 16,406 |
| GO | 12,414 | 14,532 |
| KO | 10,838 | 12,605 |
| Pfam | 20,359 | 23,651 |

**Figure 3.**
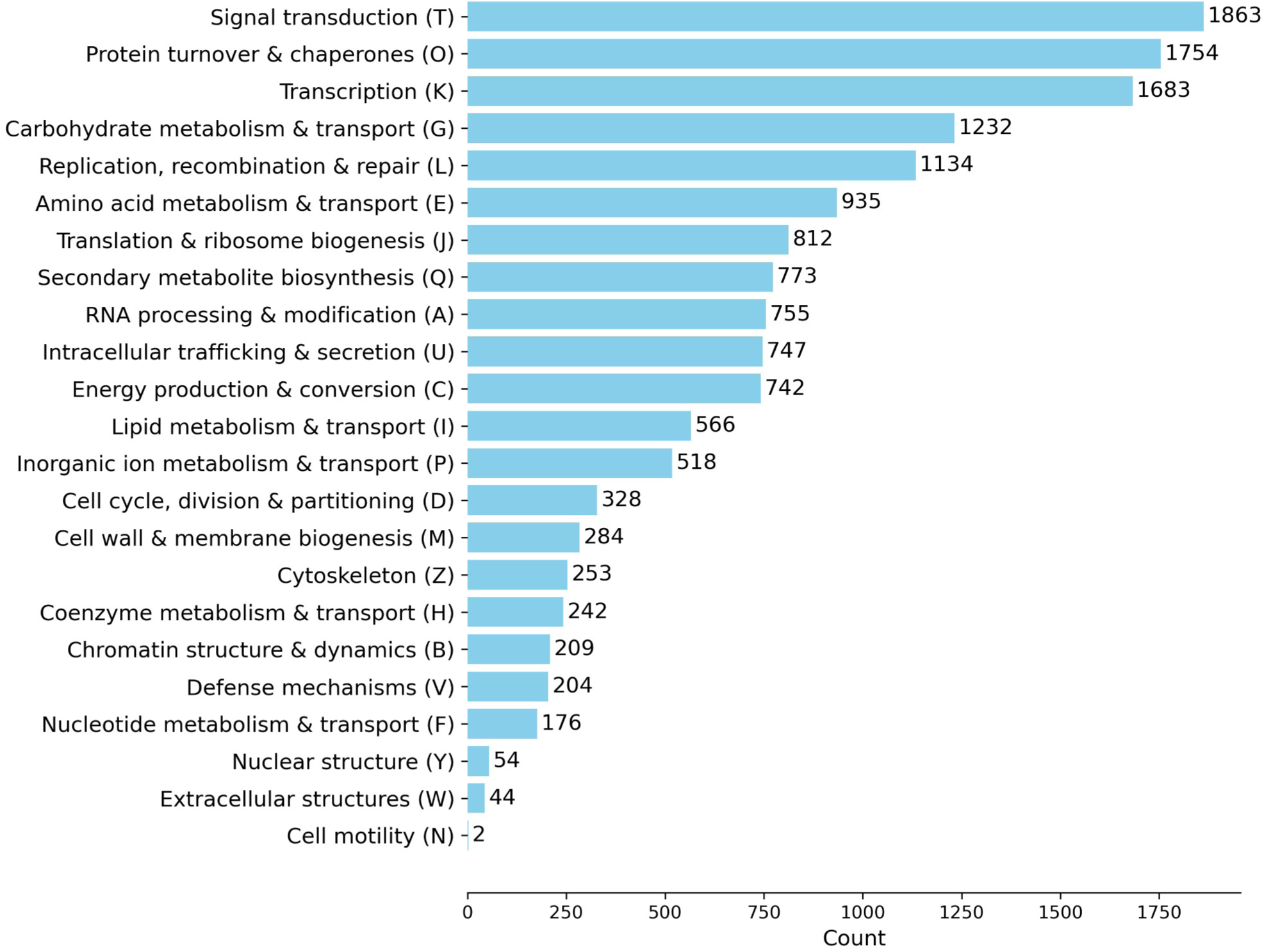
Functional annotation and category distribution of TabSet-1 genes. Distribution of annotated genes across eggNOG functional categories, showing that all major functional classes are represented in TabSet-v1.

To assess gene set completeness, BUSCO analyses were performed using both the eukaryota_odb12 and eudicotyledons_odb12 datasets. The broad eukaryota_odb12 lineage analysis detected all BUSCO genes in the annotation (100%). Using the more specific eudicotyledons_odb12 dataset, 93.6% of BUSCO genes were identified in single copy. In total, 98.1% of BUSCO genes were recovered as complete, indicating that the annotated gene set provides a highly comprehensive representation of the table beet genome (Table 5).

**Table 5.** Assessment of gene set completeness of TabSet-1. BUSCO analyses were performed with gene sets from both the general eukaryota_odb12 lineage and the more specific eudicotyledons_odb12 lineage.

| BUSCOs | eukaryota_odb12 | eudicotyledons_odb12 |
| --- | --- | --- |
| Complete | 129 (100%) | 2,752 (98.1%) |
| Single-copy | 110 (85.3%) | 2,626 (93.6%) |
| Duplicated | 19 (14.7%) | 126 (4.5%) |
| Fragmented | 0 | 18 (0.6%) |
| Missing | 0 | 35 (1.2%) |
| Total | 129 | 2,805 |

Several QTLs associated with geosmin concentration have previously been reported on chromosome 8 [6]. Within TabSet-1, three adjacent genes on BvChr 8 (tab_g19796, tab_g19798, and tab_g19804 in the region of 15.21–15.43 Mbp) encode proteins with conserved terpene synthase domains. One associated marker is located at EL10.2 position 15.30 Mbp (15.41 Mbp in the table beet assembly), placing it close to this terpene synthase gene cluster and suggesting that these genes are potential candidates for further investigation of geosmin biosynthesis.

To support the functional annotation of these TPS candidates, we performed a phylogenetic analysis including candidate TPS protein sequences from sugar beet [12], *Arabidopsis* [42], rice [43], tomato [44], and grapevine [45], together with the three proposed TPS candidates from table beet BvChr 8 (Supplementary Fig. S2). The three table beet candidates clustered with the corresponding sugar beet TPS protein sequences, whereas TPS candidates from other species formed their own subtrees. No amino acid differences were detected between the TPS candidates from table beet and sugar beet. Thus, the identified TPS candidate genes appear to be conserved between table beet and sugar beet, consistent with reports that sugar beet can also exhibit high geosmin levels [46]. However, functional validation will be required to confirm their direct involvement in geosmin biosynthesis.

In addition to protein-coding genes, we annotated non-coding RNAs (ncRNAs) using the infernal cmscan pipeline and identified 10,405 loci, of which 10,035 resided on the chromosomally assigned scaffolds. The most abundant ncRNA class was 5S ribosomal RNA (rRNA) with 7,080 occurrences, of which a large cluster encompassing 7,028 loci was located on BvChr 4 between 46 Mbp and 50 Mbp (7,037 loci in total on BvChr 4). We searched for this ncRNA class in sugar beet and found a similar clustering of 5S rRNA on BvChr 4 in the assemblies RefBeet 3.0 and FC309 hap1. The second and third most abundant classes were small nucleolar RNAs (snoRNAs; 1,203 loci) and transfer RNAs (tRNAs; 1,043 loci), both of which were distributed across the nine table beet chromosomes (Table S3).

A subsequent, more specific analysis of tRNAs predicted a total of 1,054 tRNA genes, of which 1,027 were located on the nine chromosome scaffolds (Table S4). We detected similar numbers in RefBeet 3.0, with 1,010 tRNAs on BvChr 1-9 and in FC309, with 1,017 tRNAs on BvChr 1-9.

### Genome-wide cytosine methylation profiling in table beet

To characterize the epigenetic landscape of table beet, enzymatic methyl (EM)-seq sequencing to generate genome-wide DNA cytosine methylation data was performed. The advantage of EM-seq is that it captures both 5-methylcytosine (5mC) and 5-hydroxymethylcytosine (5hmC); since 5hmC is rare or absent in plants, the signal is assumed to predominantly reflect 5mC. After quality filtering, 152.6 million read pairs (120.8x coverage of the table beet genome) were retained and aligned to RefTab-1, followed by methylation calling using the BSBolt pipeline. Global cytosine methylation levels were calculated as weighted methylation rates (total methylated counts divided by total coverage counts) in the CG, CHG, and CHH contexts, where “H” denotes any base except “G”, showing overall methylation rates of 80.7%, 49.3%, and 11.5%, respectively.

At the chromosome scale, methylation levels varied along the chromosomes and broadly corresponded to genomic feature density, with higher methylation in repeat-rich regions and lower methylation in exon-rich regions (Fig. 4A; Supplementary Fig. S3).

**Figure 4.**
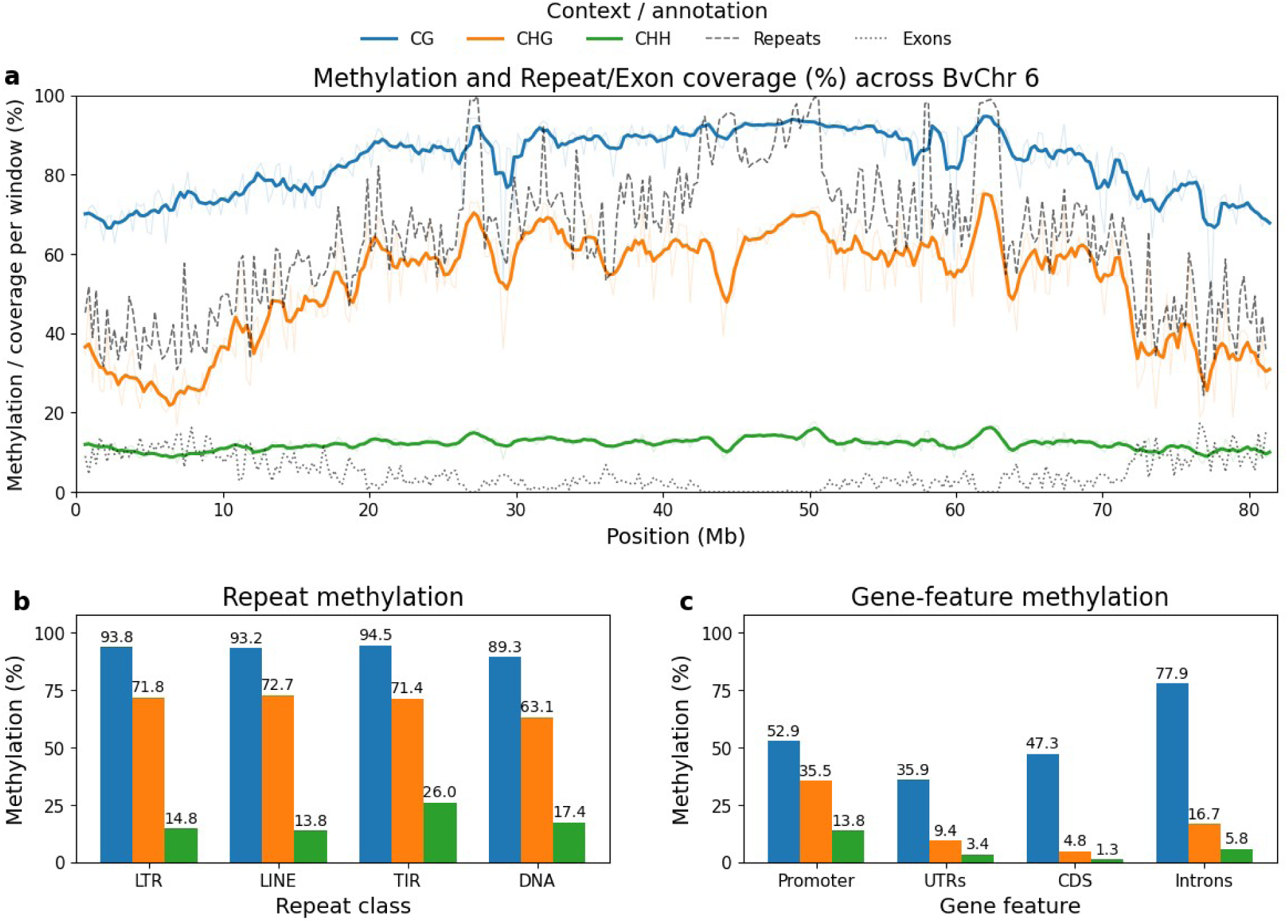
Distribution of methylation and annotations across chromosome 6 and genome-wide methylation across selected genomic feature classes. Cytosine methylation levels are shown separately for CG, CHG, and CHH sequence contexts, along with repeat and exon coverage in genomic windows of size 250 kbp (a). Average methylation levels are compared for (b) major repeat classes (LTR: long terminal repeats, LINE: long interspersed nuclear elements, TIR: terminal inverted repeats, DNA: DNA transposon repeats) and (c) gene-associated features (UTR: untranslated region, CDS: coding sequences).

We next examined methylation distributions across major genomic features. Repeat sequences displayed the highest methylation levels, consistent with the role of DNA methylation in transposon silencing. Non-repetitive intergenic regions and gene bodies showed similar methylation levels in the CG context (∼61–65%), but differed in non-CG contexts: intergenic regions displayed notably higher CHG (∼42%) and CHH (∼8%) methylation compared to gene bodies (CHG ∼12%, CHH ∼4.5%) (Table 6).

**Table 6.** Cytosine methylation levels across genomic features and repetitive elements in the RefTab-1 assembly. Values represent the percentage of methylated cytosines in the CG, CHG, and CHH sequence contexts (where H = A, C, or T). Genomic features include gene-associated regions, non-repetitive intergenic regions, and repetitive element classes, including LTR retrotransposons (LTRs), long interspersed nuclear elements (LINEs), terminal inverted repeats (TIRs), and DNA transposons. Promoters are defined as the 1,000 bp region upstream of the transcription start site.

| Genomic Feature | CG | CHG | CHH |
| --- | --- | --- | --- |
| Region |  |  |  |
| Repeat sequences | 92.90 | 70.18 | 16.04 |
| Gene bodies | 65.22 | 12.27 | 4.50 |
| Non-repetitive intergenic | 61.54 | 41.77 | 8.22 |
| Gene bodies |  |  |  |
| Introns | 77.93 | 16.72 | 5.78 |
| Exons | 46.42 | 5.12 | 1.52 |
| Coding sequence | 47.28 | 4.8 | 1.31 |
| UTR | 35.87 | 9.42 | 3.44 |
| Promoter | 52.87 | 35.51 | 13.83 |
| Repeat sequences |  |  |  |
| LTR | 93.79 | 71.75 | 14.75 |
| LINE | 93.22 | 72.71 | 13.84 |
| TIR | 94.54 | 71.37 | 25.98 |
| DNA | 89.34 | 63.13 | 17.45 |
| Total | 80.69 | 49.30 | 11.48 |

For gene-associated regions, we further analyzed the methylation status of promoter regions, exons, and introns. Promoters were defined as the 1,000 bp region upstream of the transcription start site (TSS), a commonly used definition in plant methylation studies to capture core regulatory elements. Methylation levels varied systematically across these features and sequence contexts. Exons exhibited the lowest methylation levels across all contexts (CG ∼46%, CHG ∼5%, CHH ∼1.5%). Introns showed high CG methylation (∼78%) but low levels of CHG (∼17%) and CHH methylation (∼6%). Promoters displayed intermediate CG methylation (∼53%) but strikingly elevated non-CG methylation compared to gene bodies, with CHG levels of ∼36% and CHH levels of ∼14% (Table 6, Fig. 4C).

To characterize methylation patterns in repetitive elements, we used the repeat classification of LTRs, LINEs, TIRs, and DNA transposons. LTRs, LINEs, and TIRs exhibited similarly high methylation levels in CG (>93%) and CHG contexts (>71%). DNA transposons showed comparable methylation levels at CG sites (∼89%) but had slightly lower levels at CHG sites (∼63%). In the CHH context, TIRs displayed the highest level of methylation (∼26%), followed by DNA transposons (∼17%). LTRs and LINEs also showed moderate levels of CHH methylation (∼14-15%) (Table 6, Fig. 4B).

### Comparative analysis of table beet and sugar beet genome assemblies

In order to assess the structural similarity and divergence between table beet and sugar beet genomes, we performed a comparison of the W357B table beet genome assembly with three publicly available sugar beet cultivars (KWS2320 [10], FC309 [14], and EL10 [12]). The total table beet assembly length of 668.6 Mbp is in the range of these sugar beet assemblies. In terms of sequence length, chromosome-level comparisons showed the closest similarity to FC309, with an average size difference of 2.2 Mbp. In contrast, the average differences from KWS2320 and EL10 were 9.7 Mbp and 8.8 Mbp, respectively (Table S5). Whole-chromosome alignments between RefTab-1 and the two sugar beet genotypes KWS2320 and FC309 revealed overall long-range collinearity (Supplementary Fig. S4). A notable structural difference of ∼5.5-6 Mbp in size detected on BvChr 6 appeared to represent a distinct feature in RefTab-1 shared with neither RefBeet-3.0 nor FC309. In RefTab-1, this region contains very few annotated genes, corresponding mainly to housekeeping genes, without any indications of unusual or lineage-specific gene content. A large inversion of approximately 9 Mbp was identified at the end of BvChr 9. This structural rearrangement was observed across all examined sugar beet assemblies, indicating that it is a shared feature rather than an assembly artifact (Fig. 5). While telomeric sequences are typically located at the physical ends of chromosomes, we observed a telomere-like repeat peak in RefTab-1 at the start of the inverted region, approximately 9 Mbp from the end of BvChr 9, which may be related to the identified rearrangement (Fig. 2B, Fig. 6). The telomeric repeat peak in RefTab-1 coincides with the start of the inverted region, although the breakpoint position differs slightly among the examined sugar beet assemblies, suggesting that this region may be structurally dynamic. The co-localization of the telomeric repeat peak with the inversion boundary further suggests a possible relationship between telomere-like repeats and the rearrangement history of this region. This interpretation is consistent with previous observations in yeast, where interstitial telomeric sequences were associated with gross chromosomal rearrangements, including inversions between internal telomeric repeats and terminal telomeres [47]. The inversion may therefore represent a real structural polymorphism within *Beta vulgaris*.

**Figure 5.**
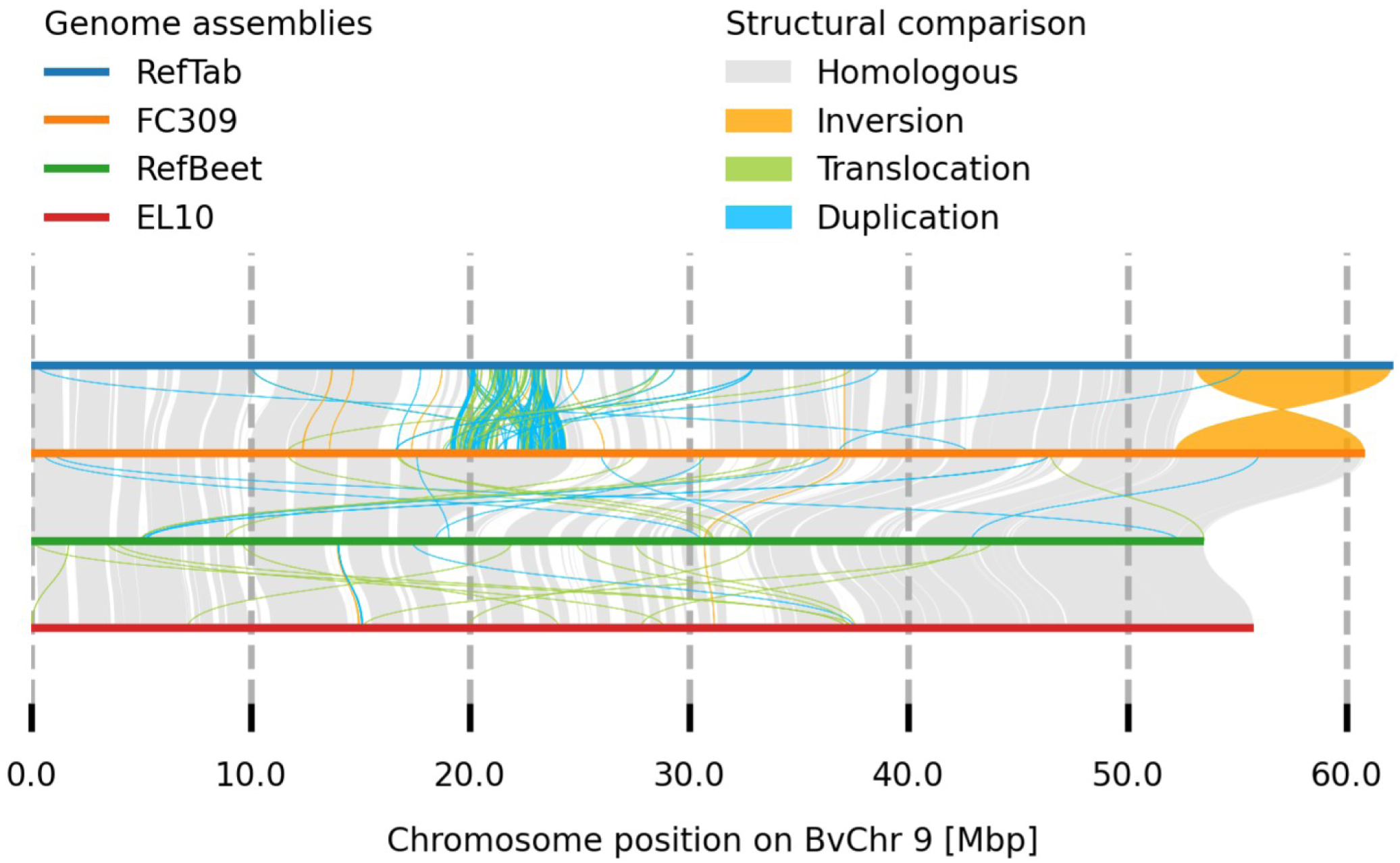
Identification of a large terminal inversion on BvChr 9. Alignment of chromosome 9 between table beet genotype W357B and sugar beet genotypes FC309, EL10, and KWS2320, showing a large inversion of approximately 9 Mbp at the end of the chromosome in W357B.

**Figure 6.**
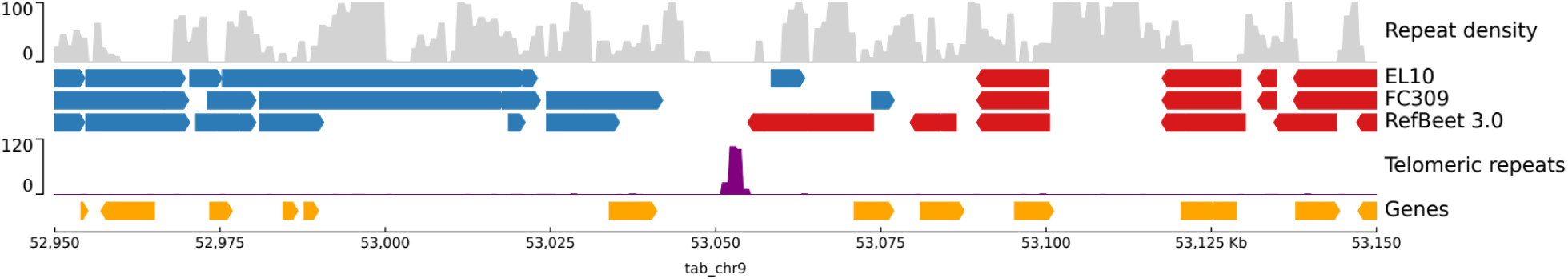
Magnified view of the internal telomeric repeat peak at the BvChr 9 inversion boundary. The region from 52.95 to 53.15 Mbp on BvChr 9 is shown together with repeat density, aligned segments from the sugar beet assemblies EL10, FC309, and RefBeet-3.0, telomeric repeat abundance, and annotated genes. Blue and red blocks indicate aligned regions in opposite orientations. The internal telomeric repeat peak in RefTab-1 occurs near the inversion boundary, while the start of the inversion differs slightly among the examined sugar beet assemblies.

## Discussion

Here we present RefTab-1, a high-quality, chromosome-scale assembly of the table beet line W357B, providing a comprehensive genomic resource for this beet cultivar type, capturing both gene-rich and repeat-rich regions. Repetitive sequences make up a large portion of RefTab-1 (63.68%), with LTR retrotransposons being the dominant repeat class. Combining multiple annotation approaches reduced the amount of unclassified repeats to 4%, demonstrating a robust repeat classification workflow.

Compared to the preliminary assembly of W357B [28], RefTab-1 represents a substantial advancement. Benefiting from a two-fold increase in PacBio HiFi read depth, the current assembly achieves superior resolution even prior to scaffolding and integrates high-confidence gene models, repeat annotations, and a genome-wide methylation profile. This enhanced structural and functional resolution is highly important for studying the crop, as previous molecular studies on table beet-specific traits, such as the identification of loci governing geosmin concentration [6], had to rely on sugar beet reference genomes. Analyzing a related but distinct taxon may obscure lineage-specific variations, such as the inversion identified on BvChr 9. A fully annotated reference genome of the quality of RefTab-1 provides an accurate genomic context for future association studies and the characterization of structural and epigenetic variation in table beet.

The nine largest RefTab-1 scaffolds, encompassing ∼96% of the total assembly size, were assigned to BvChr1–9 based on sequence homology to sugar beet. Telomeric repeats were detected at almost all (16 out of 18) ends of these scaffolds, demonstrating that the scaffolds indeed span entire chromosomes in the correct order and orientation, indicating telomere-to-telomere completeness except for two chromosomes (BvChr 1 and BvChr 3). After curation, only a small number of scaffolds remained unassigned (361 scaffolds, representing 25.3 Mbp or ∼4% of the assembly).

Comparative analysis with different sugar beet assemblies showed the closest structural similarity to sugar beet line FC309 [14], which was generated using sequencing technologies similar to those used for RefTab-1 (PacBio HiFi and Omni-C scaffolding). Differences from RefBeet-3.0 [10] and EL10.2 [12] assemblies, which relied on PacBio Continuous Long Reads (CLR) instead of Circular Consensus Sequencing (CCS), also known as HiFi, could reflect technical rather than biological differences. However, the inversion on BvChr 9 appears to be a shared feature across genomes rather than an assembly artifact, a conclusion further supported by the observation of telomeric sequences at the boundary of the inverted region in RefTab-1. The presence of a large inversion was previously mentioned within the community of sugar beet breeders and geneticists and can now be confirmed based on chromosome-wide sequence comparisons across several genotypes.

For gene prediction, we relied on sugar beet transcriptome data alongside protein homology evidence from OrthoDB [48]. While the transcriptome data may underrepresent table beet-specific genes or lowly expressed transcripts, the total number of predicted protein-coding genes (24,863) falls within the expected range for beet genomes, being slightly higher than in the EL10 assembly (24,255 genes) and lower than in RefBeet (28,271 genes in BeetSet-3). Homology-based functional annotation was achieved for 84% of the genes in TabSet-1, which was less than in BeetSet-3 (91%). Notably, three adjacent terpene synthase genes were annotated on table beet chromosome 8 in close proximity to a marker previously associated with geosmin concentration in table beet [6]. As geosmin is a sesquiterpenoid compound derived from the terpenoid pathway, we propose these genes as candidates for future investigation of its genetic basis. The number of ncRNAs in RefTab-1 (10,405) was higher than in RefBeet-3.0 (6,602). This difference was mainly due to the size of the 5S rDNA cluster on BvChr 4. In RefBeet-3.0, there were 499 annotated 5S rRNA genes on BvChr 4; in RefTab-1 this number was 7,037, while FC309 contained a total of 3,971 5S rRNA genes on BvChr 4, indicating that cluster sizes seem to vary considerably across genome assemblies and beet genotypes. Intra- and interspecific variation in rDNA loci and copy number has also been described in wheat, although accurate characterization remains challenging due to difficulties in assembling long tandem repeat arrays [49].

In the context of methylation profiling, we used Illumina short-read EM-seq data to assess the level of cytosine methylation in the genome of table beet. Although EM-seq cannot distinguish between 5mC and 5hmC, overall levels of 5hmC are very low in plant genomes [50], suggesting a negligible impact on the global profile. In addition, EM-seq avoids the DNA degradation associated with bisulfite treatment and enables reliable genome-wide methylation profiling [51]. The methylation data analyzed were from leaf tissue only (same sample as for the genome assembly), which may not capture tissue-specific or developmental stage-dependent methylation patterns. Previous comparisons in sugar beet have shown that callus tissue displays distinct hyper- and hypomethylation patterns compared to leaves, particularly in repetitive elements [38]. Thus, the methylation landscape based on other tissues may deviate from the picture obtained in this study. Also, the obtained methylation profiles may be less reliable in highly repetitive regions, because the prerequisite for exact read mapping is that reads map uniquely, a condition that is less likely to be met in repetitive sequences. This is a notable challenge in *B. vulgaris*, which has been characterized as a “methylation outlier” among angiosperms, displaying the highest global methylation levels (especially in CHH contexts) among 34 species [39]. While PacBio HiFi kinetics could, in principle, allow direct methylation detection, current models are trained on human data and therefore only support the CG context, limiting this approach for the analysis of plant genomes, which additionally carry methylation marks in CHG and CHH motifs. Despite these challenges, our analysis of specific repeat classes mirrors previously observed trends in sugar beet, where retrotransposons are heavily methylated in symmetric contexts (CG/CHG), while DNA transposons are particularly enriched for asymmetric CHH methylation [38]. We observed a similar pattern in table beet, with TIRs and DNA transposons showing the highest CHH levels (∼17–26%). However, unlike some plant genomes (e.g., Poaceae) where LTR retrotransposons are depleted of CHH methylation [39], we found moderate CHH levels (∼14–15%) in LTRs and LINEs.

## Methods

This study aimed to generate and characterize a high-quality reference genome assembly for the inbred table beet genotype W357B. It was designed as a genomics and bioinformatics study integrating long-read sequencing, Omni-C, RNA-seq, and EM-seq data for genome assembly, scaffolding, annotation, repeat analysis and methylation profiling. Leaf tissue from the inbred table beet genotype W357B was used as biological material. Experimental and computational work was performed using a combination of institutional resources and external service providers. Analyses were primarily descriptive and computational; no formal statistical hypothesis testing or power calculation was applied.

### DNA collection and sequencing

Genomic DNA was extracted from leaf tissue of the inbred table beet genotype W357B [40] using a NucleoSpin Plant II kit (Macherey-Nagel, Düren, Germany). The preparation of Pacific Biosciences (PacBio) sequencing libraries using SMRTbell adapters was outsourced to service providers, where PacBio sequencing on the Sequel II using chemistry that supports the production of HiFi data was performed. Two datasets were generated, one at USDA-ARS (NCBI SRA: SRX17632321) and one at BOKU (NCBI SRA: SRR37504297). Omni-C reads were obtained by contracting Dovetail Genomics (NCBI SRA: SRX17632322). For the genome assembly, all three datasets were combined. For cytosine methylation analysis, EM-seq data (NCBI SRA: SRR37504296) were generated from genomic leaf DNA on an Illumina NovaSeq instrument. BOKU contracted library preparation and sequencing using PacBio and Illumina technologies through Vienna Biocenter Core Facilities (VBCF).

### Preparation of sequencing data

Circular consensus (CCS) reads were generated from PacBio subreads using ccs v6.4.0 [52], and HiFi reads were extracted with extracthifi v3.5.0 [53]. The reads were converted to FASTQ format using bam2fastq v3.5.0 [53]. Reads were polished using the DeepConsensus v1.2.0 pipeline [54] with parameter “--min-rq=0.88” for ccs and the model set to “checkpoint-1”. Omni-C reads were bridge-trimmed using cutadapt v4.5 [55] with parameters “-b GGTTCGTCCA -B GGTTCGTCCA”. Quality trimming was performed using trim_galore v0.6.10 [56] with parameter “--paired” for both Omni-C reads and EM-seq reads. The success of quality trimming was checked using fastqc v0.12.1 [57] with parameter “-t 10” for increased memory.

### Genome assembly, quality assessment, assembly comparisons

HiFi reads were assembled using hifiasm v0.19.8 [58] with the parameters “-a 4 -l 3” for rounds of assembly cleaning and duplication purging. The graphical fragment assembly (GFA) file was converted to FASTA format using awk (GNU Awk 4.2.1) with the command “awk ’/S/{print “>”$2;print $3}’”. For scaffolding, quality-trimmed Omni-C reads were mapped to the table beet assembly using bwa v0.7.19 [59] with parameter “-5SP”. Duplicates were marked using samblaster v0.1.26 [60] and filtering was done using samtools v1.20 [61] with parameters “-S -h -b -F 3340”. Another filtering step was performed using the “filter_bam” script from HapHiC [62] v1.0.5 with parameters „1 --nm 3”. Scaffolding was performed using yahs v1.2a [63] with parameter “-r 10000,20000,50000,100000,200000,500000,1000000,2000000,5000000,10000000,20000000,500 00000,100000000,200000000,500000000”. Assembly contamination was scanned using FCS-GX from the NCBI Foreign Contamination Screen tool suite v0.5.4 [64] with parameter “--tax-id 3555”, corresponding to *Beta vulgaris* subsp*. vulgaris*. BLAST v2.16.0 [65] with the ncbi core_nt database and parameter “-outfmt 6” was used to search for organelle-derived contigs. These contigs were removed from the assembly using seqkit v2.10.1 [66] with subcommand grep and parameters “-v -n”. Minimap2 v2.30 [67] with parameter “-x asm5” was used to check for duplicated contigs, followed by removal of contigs with a coverage >99% in the chromosome scaffolds. The chloroplast genome was assembled using TIPPo v1.3.0 [68] with parameter “-g chloroplast”. The resulting genome was then compared with the sugar beet chloroplast genome (NCBI: NC_059012.1) using blastn v2.16.0 with parameter “-outfmt 6”.

BUSCO v5.8.3 [69] was used for completeness estimation with parameters “--miniprot -m genome” and parameter “--lineage_dataset” set to “eudicotyledons_odb12” or “eukaryota_odb12”. Telomere repeat regions were searched for with tidk v0.2.65 [70] and parameter “-c Caryophyllales”.

Alignments between the genome assemblies of table beet and different sugar beet genotypes were done using minimap2 v2.30 with parameters “-x asm5 -c “. Size differences between chromosomes were calculated based only on the size of the pseudochromosome scaffolds; no alignment lengths were taken into account.

### Repeat modeling and masking

RepeatModeler [71] and RepeatMasker [72] were used via Dfam TETools v1.88 [73]. RepeatModeler was used with parameters “-LTRStruct -genomeSampleSizeMax 750000000”. Repeat classification was done using Repbase library v20181026 [74] and the Dfam database [75]. RepeatMasker was used with parameter “-lib” set to the newly generated library from RepeatModeler. The repeat annotation was further used to mask repeats in the genome assembly as a prerequisite for gene annotation by using bedtools v2.27.1 [76] with subcommand “maskfasta” and parameter “-soft”. To provide a general repeat annotation, EDTA v2.2.2 [77] was run with parameters “--curatedlib” pointing to the BeetRepeats repeat library [78], “--rmlib” pointing to the RepeatModeler repeat library, “--cds” pointing to the coding sequences from gene prediction, “-- exclude” pointing to a BED file with gene coordinates and “--anno 1”.

### Gene annotation

Approximately 2.15 billion read pairs and 408 million single-end reads from mRNA-seq (NCBI SRA: ERR11534264-ERR11534303, ERR11534320, ERR11534330-ERR11534345, SRR1508751, SRR1508753, SRR1508755, SRR1508756, SRR1508758, SRR868886, SRR868887, SRR869751-SRR869758) were mapped to the RefTab-1 assembly using hisat2 v2.2.1 [79] with parameters “--phred33 --max-intronlen 20000 --dta” and sorted using samtools v1.21 with subcommand “sort”. BRAKER3 v3.0.7.5 [80,81] with parameter “--genome” with the softmasked genome, “--prot_seq” with the Viridiplantae dataset from OrthoDB 11 [48], “--bam” with the mapped mRNA reads, “--species=beta_vulgaris” which is an Augustus [82] species model generated previously [18] and “--augustus_args=’--noInFrameStop=true --genemodel=complete’ -- useexisting --busco_lineage=eudicots_odb10”. StringTie [83], run with default parameters, and a Python script were used to add UTR information to the gene annotation. Functional annotation was performed using eggNOG-mapper v2.1.12 [84,85] with parameters “-m diamond --itype CDS -- go_evidence all”, using DIAMOND for sequence searches [86]. Gene Ontology (GO) terms [87], KEGG annotations [88], Pfam protein families and domains [89], and COG functional categories [90] were obtained from the eggNOG-mapper output. BUSCO v5.8.3 was used for completeness estimation with parameter “--miniprot -m prot” and parameter “--lineage_dataset” set to “eudicotyledons_odb12” or “eukaryota_odb12” respectively. To calculate the TPS phylogeny we first annotated the gene sets of the different species using eggNOG-mapper v2.1.12 with parameters “-m diamond --itype CDS --go_evidence all”. Genes with a Pfam annotation containing “Terpene_synth” or “Terpene_synth_C” and a minimum length of 400 amino acids were extracted. For genes with multiple transcripts, only the longest transcript was retained. Duplicate genes within a species were removed using “seqkit rmdup”. MAFFT v7.453 [91] with parameter “--auto” was used to align the sequences, followed by trimAl v1.5.rev1 [92] with parameter “--automated1” for automated alignment trimming. IQ-TREE v2.4.0 [93] with parameters “-m MFP -B 1000 -alrt 1000 -T AUTO” was used to calculate the phylogenetic tree. Infernal cmscan v1.1.4 [94] with the Rfam covariance models release 14.5 [95] and parameters “--cut_ga --rfam --nohmmonly --oskip --fmt 2” was used for annotation of RNA families. We used tRNAscan-SE v2.0.12 [96] with parameter “-E” for tRNA prediction.

### Methylation annotation

BSBolt v1.6.0 [97] was used for methylation annotation. An index of the RefTab-1 genome assembly was created using subcommand “Index”, then reads were aligned using subcommand “Align”. Samtools v1.21 was used for methylation calling preprocessing with subcommand “fixmate” and parameter “-p” and subcommand “markdup”. Lastly BSBolt subcommand “CallMethylation” with parameter “-verbose” was used for calling methylation.

### Scripting, plotting and data visualization

Python 3 [98], R 4.5.1 [99] and Perl 5 [100] were used for scripting, plotting, and data visualization. Python libraries used were matplotlib v3.8.1 [101], numpy v1.26.0 [102], pandas v2.1.3 [103] and pyranges v0.1.4 [104]. Circlize v0.4.16 [105] was used as plotting library in R. Additionally, jbrowse2 v3.6.4 [106], IGV v2.19.5 [107], syri v1.7.1 [108], plotsr v1.1.1 [109], iTOL [110] and pyGenomeTracks v3.9 [111] were used for data visualization.

### Genome assemblies and datasets analyzed

The publicly available sugar beet genome assemblies used for comparative analyses were obtained from the following sources: RefBeet-3.0 (genotype KWS2320) from https://bvseq.boku.ac.at/Genome/Download/RefBeet-3.0/, EL10.2 from NCBI Genome under accession GCA_026745355.1, and FC309 hap1 from NCBI Genome under accession GCA_037177695.1.

The publicly available gene annotations used to build the phylogenetic tree of TPS genes were obtained from the following sources: sugar beet from NCBI Genome under accession GCA_026745355.1, *Arabidopsis* from https://www.arabidopsis.org/download/list? dir=Genes%2FAraport11_genome_release, rice from https://rice.uga.edu/download_osa1r7.shtml, tomato from https://solgenomics.net/ftp/tomato_genome/annotation/ITAG4.0_release/, and grapevine from https://grapedia.org/files-download/.

### Computing resources and data management

Data were analyzed using a Linux cluster running CentOS 7 and AlmaLinux 8 on nodes with up to 64 cores and at most 1 TB RAM. Additionally, analyses were performed on VSC-5 nodes of the Austrian Scientific Computing infrastructure using up to 128 cores and 1 TB RAM. Alongside the computing infrastructure, a range of command-line utilities and workflow tools were employed to support data processing, sequence handling, and parallelization. Among the tools used were: parallel v20190622 [112], bioawk v20110810 [113], conda-forge [114], bioconda [115], miniforge [116] and standard Unix text processing utilities (sed, awk, grep, cut). Format conversions, calculations of assembly metrics, and methylation analysis, including the assessment of methylation levels and percentages across different genomic regions (genes, repeats, promoters, and intergenic regions) were performed with standard Linux tools and scripts written in Perl and Python.

## Availability of data and materials

The datasets generated and analyzed during the current study are available in the NCBI Sequence Read Archive (SRA) under accession numbers SRR37504297 and SRX17632321 (PacBio data), SRX17632322 (Omni-C data), and SRR37504296 (EM-seq data). The genome assembly (RefTab-1), gene models (TabSet-1), further annotations, and methylation data are available at https://bvseq.boku.ac.at/Genome/Download/. Python scripts used for gene annotation post-processing and methylation analysis in this study are available via Zenodo at https://doi.org/10.5281/zenodo.22278055.

## Authors’ contributions

H.H., K.D., and J.C.D. conceived the study. T.H. performed analyses and drafted the manuscript. L.A.R. performed initial analyses. J.C.D. and H.H. revised the manuscript including the comments of all authors.

## Acknowledgements

We are grateful to Irwin Goldman for providing seeds of table beet line W357B, Christian Gottschall for technical support, Nancy Stralis-Pavese for DNA extraction, and Lukas Schönmann for providing the picture of a table beet. The computational results were in part achieved using the Austrian Scientific Computing infrastructure. Genomic sequencing was in part performed by the Next Generation Sequencing Facility at Vienna BioCenter Core Facilities (VBCF), a member of the Vienna BioCenter (VBC), Austria.

## Supplementary table legends

**Table S1:** Genome-wide repeat annotation of the RefTab-1 (W357B) assembly using the Extensive de novo TE Annotator (EDTA). The analysis combined EDTA with the RepeatModeler output and the BeetRepeats library. The table shows counts, masked bases, and percentages for each repeat class. Major repeat groups include long interspersed nuclear elements (LINEs), long terminal repeats (LTRs), terminal inverted repeats (TIRs), and ribosomal DNA (rDNA). Other classes include short interspersed nuclear elements (SINEs), non-LTR retrotransposons, non-TIR DNA transposons, satellite DNA, and fragmented repeats.

**Table S2:** Summary statistics of TabSet-1. Counts and size metrics for genes, transcripts, exons, introns, and untranslated regions, including averages, total lengths, and maximum lengths.

**Table S3:** Summary of annotated non-coding RNAs (ncRNAs) in the RefTab-1 assembly. Counts are shown per ncRNA class as identified by the Infernal cmscan pipeline. The most abundant category is ribosomal RNA (rRNA), followed by small nucleolar RNAs (snoRNAs) and transfer RNAs (tRNAs).

**Table S4:** Predicted tRNA genes in the RefTab-1 assembly. Counts of tRNA genes per amino acid, as identified by tRNAscan-SE, are shown, grouped by amino acid class (nonpolar, polar uncharged, acidic, basic) and special categories (initiator methionine, suppressor codon, undetermined).

**Table S5**: Chromosome assembly sizes in assemblies of different *Beta vulgaris* genotypes, including table beet and sugar beet. Genotypes and assembly versions: table beet W357B (RefTab-1); sugar beets KWS2320 (RefBeet-3.0), FC309 (FC309 hap1), and EL10 (EL10.2). Avg. Diff.: Size difference in Mbp averaged over all chromosomes (as absolute values).

